# Transcriptomic Changes During Stage Progression of Mycosis Fungoides

**DOI:** 10.1101/2021.04.20.440708

**Authors:** M Xiao, D Hennessey, A Iyer, S O’Keefe, F Zhang, A Sivanand, R Gniadecki

## Abstract

**Background:** Mycosis fungoides (MF) is the most common cutaneous T cell lymphoma, which in the early patch/plaque stages runs an indolent course. However, ~25% of MF patients develop skin tumors, a hallmark of progression to the advanced stage and associated with high mortality. The mechanisms involved in stage progression are poorly elucidated.

**Methods:** We performed whole-transcriptome and whole-exome sequencing of malignant MF cells from skin biopsies obtained by laser-capture microdissection. We compared three types of MF lesions: early-stage plaques (ESP, n=12), and plaques and tumors from patients in late-stage disease (late-stage plaques, LSP, n=10, and tumors, TMR, n=15). Gene Ontology (GO) and KEGG analysis were used to determine pathway changes specific for different lesions which we linked to the recurrent somatic mutations overrepresented in MF tumors.

**Results:** The key upregulated pathways during stage progression were those related to cell proliferation and survival (MEK/ERK, Akt-mTOR), Th2/Th9 signaling (IL4, STAT3, STAT5, STAT6), meiomitosis (CT45A1, CT45A3, STAG3, GTSF1, and REC8) and DNA repair (PARP1, MYCN, OGG1). Principal coordinate clustering of the transcriptome revealed extensive gene expression differences between early (ESP) and advanced-stage lesions (LSP and TMR). LSP and TMR showed remarkable similarities at the level of the transcriptome, which we interpreted as evidence of cell percolation between lesions via hematogenous self-seeding.

**Conclusion:** Stage progression in MF is associated with Th2/Th9 polarization of malignant cells, activation of proliferation, survival, as well as increased genomic instability. Global transcriptomic changes in multiple lesions are probably caused by hematogenous cell percolation between discrete skin lesions.

## Introduction

Mycosis fungoides (MF) is the most prevalent form of primary cutaneous T-cell lymphoma (CTCL), a heterogeneous group of extranodal lymphoproliferative disorders that involve the skin.^1^ Early-stage (IA-IIA) disease is characterized by erythematous papules, patches, and plaques, and the majority of patients (80-90%) experience an indolent disease course with slow progression over years or even decades.^2–5^ However, ~25% of MF patients progress to advanced-stage (IIB-IV) disease, distinguished by the appearance of skin tumors, extracutaneous dissemination to lymph nodes, blood, and viscera, as well as a dramatic reduction in survival (decrease in 5-year survival from approximately 80% to 26%).^2–5^ The molecular mechanisms underpinning stage progression in MF remain largely unknown. Moreover, it has not been explained how the emergence of even a single tumor in a relatively small area of the skin impacts the global evolution and prognosis of MF. It would be logical to propose that the more aggressive cancer cells in skin tumors spread hematogenously to other skin areas, increasing tumor load and the clinical impact of the malignancy. However, the putative hematogenous percolation of tumor cells between different lesions has not been investigated because MF cells have been considered to represent mutated tissue-resident T-cells which tend to remain arrested in the defined areas of the skin and do not recirculate via lymphatics and the lymph nodes like the transient passenger T-cells.^6^ We reasoned that the hypothesis that MF cells traffic between skin lesions can be addressed by comparing transcriptomic profiles of skin lesions in different clinical stages of MF.

Previous analysis of the mutational landscape of CTCL revealed the emergence of numerous subclones and very complex patterns of clonal driver mutations, with little consistency among patients.^7–11^ Over the past decade, studies have focused on the analysis of the transcriptome in an attempt to characterize cancer driver genes and signaling pathways, but the results have been discordant.^12^ This is in part due to the high degree of intratumoral heterogeneity as well as the lack of standard approaches in both sample selection and experimental design that hinders direct comparison between different transcriptome studies.^12^ Many studies pooled samples from MF and the leukemic CTCL (Sézary syndrome, SS) in spite of the data indicating that these entities may represent separate diseases^6,13^ and few transcriptome studies account for the stage of disease and the morphology of skin lesions (e.g., patch, plaque, or tumor). Moreover, the samples used for transcriptomics were usually crude skin biopsies, in which the proportion of neoplastic cells is quite low, often below 30%.^7^

In this study, we compared transcriptome profiles of early and advanced MF to reveal potential mechanisms of disease progression. To capture stage-dependent expression changes, we classified MF skin lesions according to the patient’s clinical disease stage (early [stage I] vs advanced [stage ≥ IIB]) and the morphology of the lesion (plaque vs tumor). To circumvent earlier limitations, we used laser capture microdissection of neoplastic cell clusters to eliminate signal from normal cells. We were able to confirm some of the previously reported pathways associated with advanced-stage MF and identified new transcriptomic changes, such as upregulation of pseudo-meiotic processes (meiomitosis) that may be involved in tumor progression. Moreover, we found that stage progression is associated with global changes in transcriptomic profiles affecting multiple MF lesions which we interpret as evidence of cancer cell percolation proposed by us previously^14^ and analogous to tumor self-seeding.^15,16^

## Methods

### Samples, whole transcriptome, and whole-exome sequencing

Ethics approval was obtained under the application HREBA.CC-19-0435 from the Institutional Review Board at the University of Alberta. 37 laser-microdissected skin lesions were collected from 23 adult patients with a diagnosis of MF in stages IA to IVB (**Supplementary Table S1**), as previously described^7^. RNA was isolated from the microdissected tissues using AllPrep DNA/RNA Micro kit from Qiagen (Hilden, Germany). For samples MF4_2, MF4_3, MF5_1, MF7_1, MF7_2, MF11, MF11_1, MF19_1, MF19_2, and MF19_3 rRNA was inhibited using NEBNext rRNA Depletion kit (NEB, Ipswich, Massachusetts, USA) and later processed using NEBNext Ultra II Directional RNA Library Prep kit for Illumina (Ipswich, Massachusetts, USA). The rest of the samples in the study were processed using Ovation Solo RNA-seq system (Nugen, Redwood city, CA, USA). The manufacturer’s instructions were followed and no modifications were made to the protocols. The RNA libraries were sequenced on an Illumina HiSeq 1500 sequencer using paired-end 150 kits (cat# PE-402-4002) (Hiseq PE rapid cluster kit V2) or NovaSeq 6000 S4 reagent kit 300 cycles (cat# 20012866). Previously published whole-exome sequencing from the same samples was used for the analysis of single nucleotide variants, as described.^17^

Skin biopsies were categorized into three lesion types according to the clinical stage and morphology of the lesion at the time of collection. Plaque lesions sampled from patients in early-stage MF (IA-IIA) were designated as ESP (early-stage plaque, n=12). Patients with advanced-stage disease (≥IIB) have two types of skin lesions, the TMR (tumors, n=15) and the plaques which either persist from the early stages (late-stage plaques, LSP, n=10). There was no significant difference in tumor cell fraction between microdissected samples from different lesion types.

### Bioinformatic analysis of gene expression data

The bioinformatics pipeline involved a series of aligners and statistical methods shown in **Figure 1.** All programs were used with default or developer-supplied settings unless otherwise specified. Code is available at https://github.com/d-henness/deseq2_project. Paired fastq files were preprocessed using fastp (v 0.19.7) with the --detect_adapter_for_pe flag enabled.^18^ The resulting fastq files were then aligned and quantified using RSEM (v. 1.3.3).^19^ RSEM was used with Bowtie 2 (v. 2.4.1)^20^ as its backend aligner and an index constructed using Ensembl’s 98 human genome annotation.^21^ This yielded gene expression levels in units of transcripts per million (TPM).

**Figure 1.**
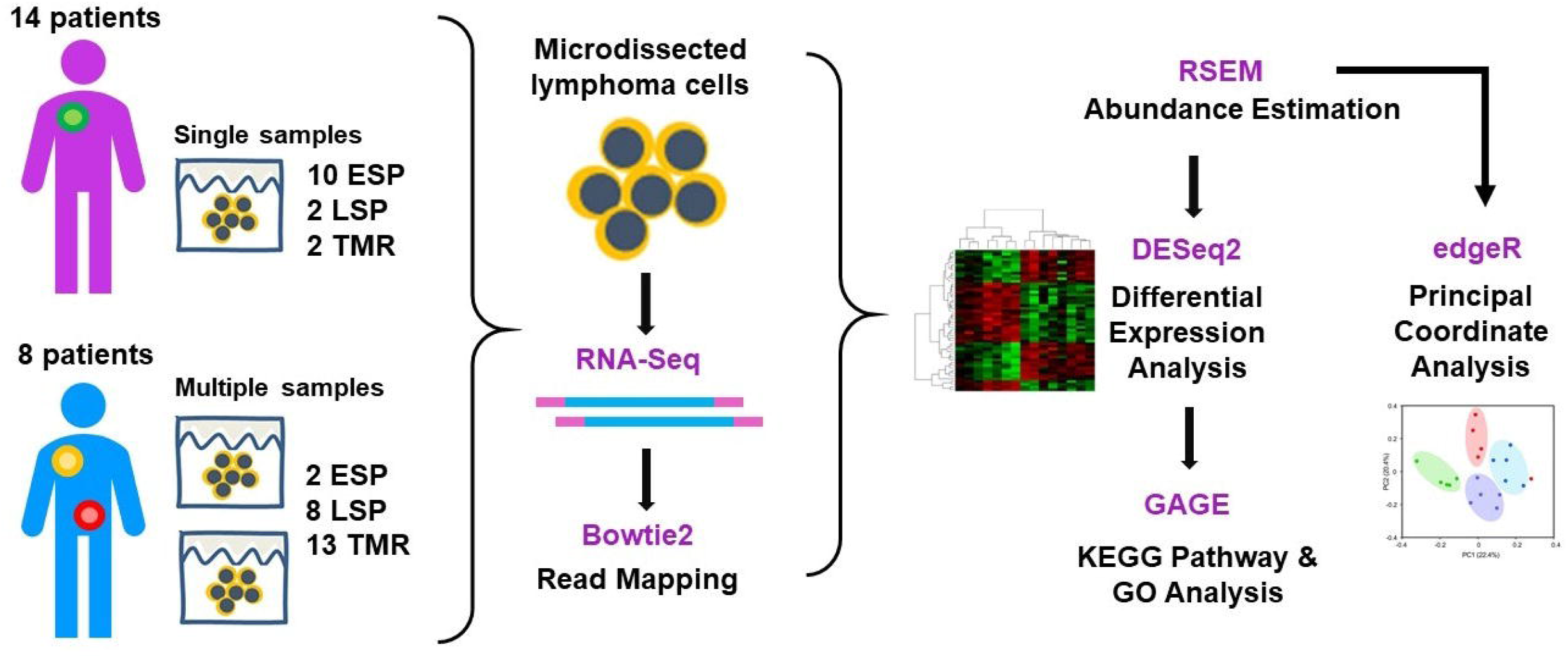
Summary of methods and study design. 32 biopsies of lesional skin were obtained from 23 patients, consisting of 14 patients that contributed single biopsies and 8 patients where multiple samples were taken. The lesions were categorized according to the both the clinical stage (advanced vs. early) as well as the morphological feature (plaque vs lesion): ESP (n=12 early stage plaque biopsies from patients in stage IA-IIA), LSP and TMR (n=10 late-stage plaques and 15 tumors biopsies from patients in stage ≥IIB, respectively). After RNA-Seq, reads were aligned to the reference sequence after removal of adapter sequences using Bowtie2. Transcript abundance was measured using RSEM. Two statistical methods were used for differential expression analysis and principal coordinate analysis (DESEq2 and edgeR, respectively). Generally applicable gene set enrichment (GAGE) analysis was performed to provide more information about the biological functions and pathways significantly enriched in each disease stage.

Unsupervised Principal Coordinates Analysis (PCoA) was performed using edgeR (R package, v. 3.30.3)^22^ to obtain an overview of the overall structure of the RNA-Seq data in a low-dimensional subspace. PCoA provides insights into the association between samples based on gene expression patterns and detects the formation of clusters among individual specimens. The DESeq2 program, implemented as a package in R (v. 1.28.1)^23^ was used to identify differentially expressed genes (DEGs) that differed significantly between pairwise disease stages (log2 fold change (log2FC)>|1|). Benjamini-Hochberg correction was used to correct for multiple comparisons, with a standard false discovery cut-off (FDR) of <0.05.

To gain further insight into the functional processes differing between lesions, we used all of the available gene expression data (cutoff-free) and performed gene set analysis using the Generally Applicable Gene Set Enrichment (GAGE) method implemented as a package in R (v. 2.37.0).^24^ We focused on Gene Ontology (GO) Biological Process (BP) terms^25^ and Kyoto Encyclopedia of Genes and Genomes (KEGG) pathways.^26^ In both databases, Benjamini-Hochberg correction was used to correct for multiple comparisons (FDR <0.05). Visual data representations were created using the R packages “ggplot2” (v. 3.3.1), “pheatmap” (v. 1.0.12), and ‘EnhancedVolcano’ (v. 1.6.0).^27^

### Data availability

The exome sequencing data is available on dbGaP under accession number phs001877.v1.p1. The RNA sequencing fastq files are available on Sequence Read Archive (SRA) under accession number....^1^

## Results

### Changes in transcriptome signatures during stage progression of MF

Principal coordinates analysis (PCoA) was used to examine the associations between our samples (12 ESP, 10 LSP, 15 TMR) and revealed three clusters (**Figure 2A**). Cluster A contained the majority of ESP samples (11/12, or 91.7%) whereas the majority of advanced-stage samples (LSP and TMR) were distributed across clusters B1 and B2 (18/25, or 72% consisting of 8/10 LSP samples and 10/15 TMR samples). This suggests the transcriptome profile of ESP is distinct from the late-stage disease whereas the expression patterns among advanced lesions (LSP and TMR) are similar. Thus, the MF transcriptomic profile reflects the stage of the disease (advanced vs early) rather than the morphology of the lesion (plaque vs tumor) and indicates that significant global alterations in gene expression occur with progression to advanced-stage disease.

**Figure 2.**
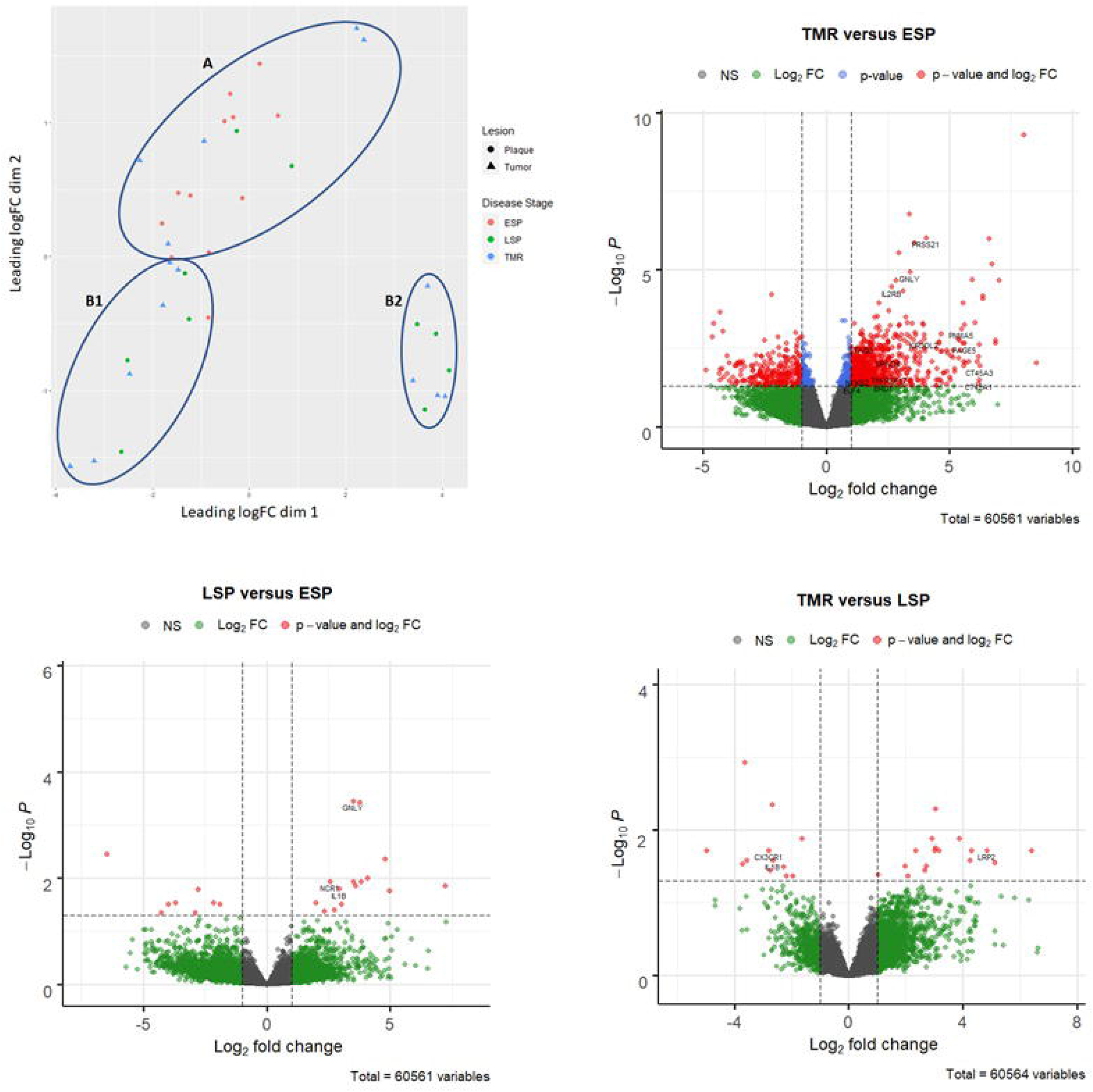
Comparative transcriptome analysis of ESP, LSP, and TMR lesions. **A.** Unsupervised Principal coordinate analysis (PoCA) showing the variation and clustering among ESP, LSP, and TMR lesion types, with ellipses drawn manually. LSP resembles TMR (clusters B1, B2), rather than ESP (cluster A). **B.** Volcano plot of differential expressed genes (DEGs) between TMR (comparison) and ESP (reference) group, generated using the EnhancedVolcano R package. The vertical y-axis) corresponds to the mean expression value of log10 (adjusted P value) after correction for multiple testing, and the horizontal x-axis displays the log2 fold change value. 1,154 transcripts with an adjusted p-value < 0.05 and |log2 (fold change)| ≥ 1 were mapped as DEGs (depicted as red dots). Genes that failed to reach statistical significance are present in green, while transcripts below fold-change cutoffs are presented in blue. Genes upregulated or downregulated in TMR compared to ESP are in the upper right and upper left quadrants, respectively. **C.** Volcano plot for the distribution of gene expression between LSP (comparison) and ESP (reference), mapping 26 DEGs (red dots). **D.** Volcano plot for the distribution of gene expression between TMR (comparison) and LSP (reference), mapping 29 DEGs (red dots).

Next, we performed pairwise comparative analyses of the expression profiles between ESP, LSP, and TMR samples to identify differentially expressed genes (DEGs) between different lesion types. Differential expression analysis of TMR vs ESP (reference) lesions identified 1,154 DEGs (fold change>2; adjusted P<0.05), of which 908 DEGs were significantly up-regulated and 246 DEGs were downregulated in TMR compared to ESP (**Figure 2B**). Fewer DEGs were detected for the comparisons of LSP to ESP (26 DEGs, 18 upregulated in LSP, **Figure 2C**) and TMR to LSP (29 DEGs, 11 upregulated in TMR, Figure 2D). A list of all differentially regulated genes across all comparisons is presented in **Supplementary Tables S2-S4**.

To further illustrate the relationship of genes found to be differentially expressed in each pairwise comparison, we used Venn diagram analysis and overlapped the 1,154 DEGs in TMR vs. ESP with the 26 DEGs in LSP vs. ESP (**Figure 3**). This revealed 1,137 DEGs unique to TMR (895 upregulated, 242 downregulated), 9 DEGs unique to LSP (5 upregulated, 4 downregulated), and a signature of 17 intersecting DEGs common to both LSP and TMR samples (13 upregulated, 4 downregulated), all concordantly altered (i.e., increased in both, or decreased in both). The large overlap in DEGs shared between LSP and TMR (both advanced lesions) compared with ESP corroborates the PCoA results and suggests that progression to advanced-stage affected the global transcriptomic profile of MF skin lesions.

**Figure 3.**
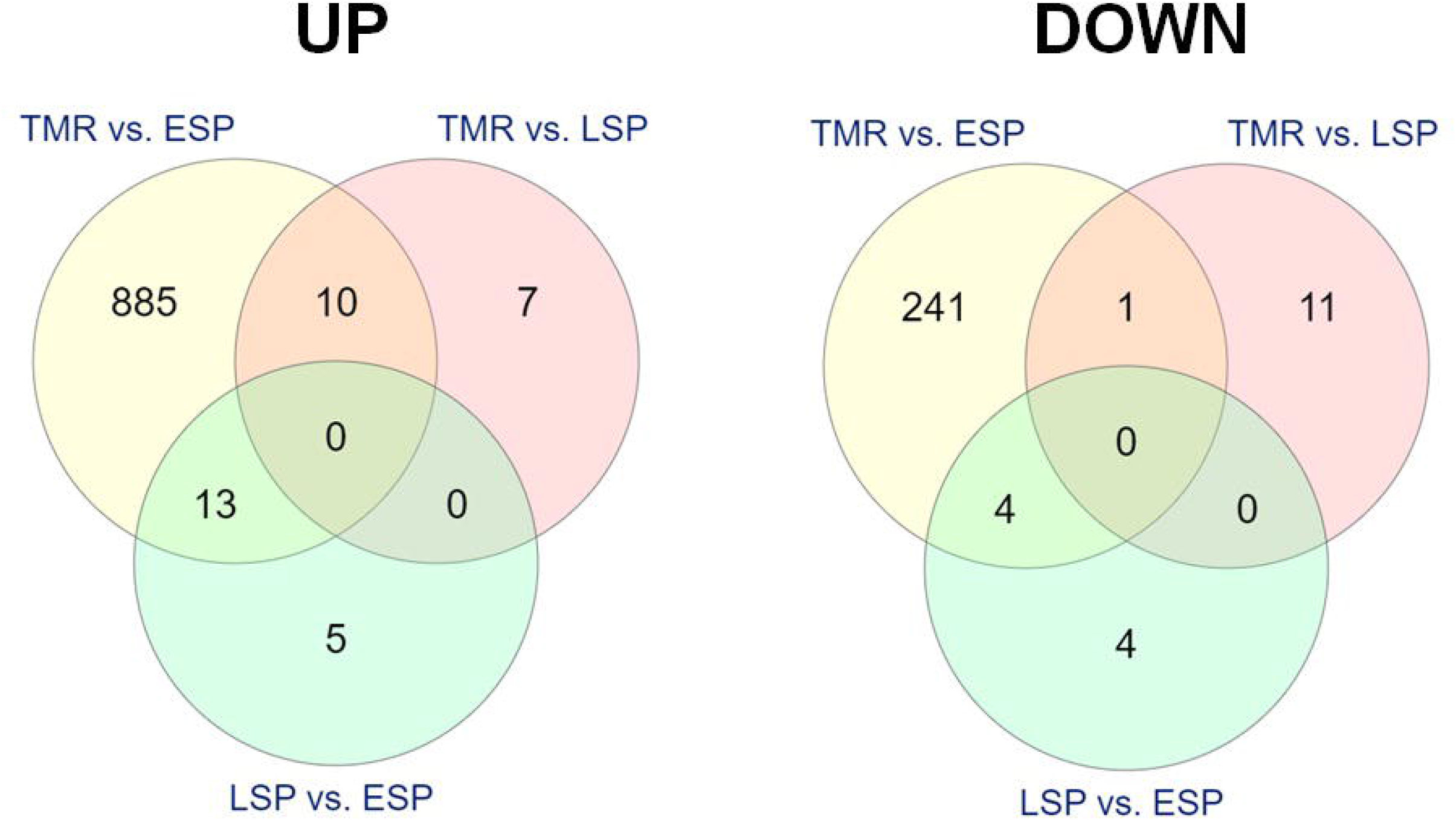
Venn diagrams analysis of differentially expressed genes. The yellow, red and green circles represent the number of up or down-regulated DEGs between each pairwise comparison (TMR vs. ESP, TMR vs. LSP, and LSP vs. ESP). The overlapping number represents mutual, concordantly changed DEGs common to each comparison and the non-overlapping number represents genes unique to each comparison.

### GO and KEGG Functional Enrichment Analysis

We performed pairwise gene set analysis (GSA) using the Gene Ontology (GO) and Kyoto Encyclopedia of Genes and Genomes (KEGG) databases to identify enriched terms and pathways at different MF stages. We used all of the available gene expression data instead of prefiltering for a list of strong DEGs in order to capture coordinated, low-level changes across genes that may belong to common pathways and are regulated by the same transcriptional network.

Our analyses identified 58 enriched GO biological processes and 21 KEGG pathways in TMR compared to ESP (**Supplementary Figure S1**). The results of GO analyses revealed that TMR lesions were significantly enriched in proliferative signaling (‘Ras protein signal transduction’, ‘activation of MAPK kinase kinase activity’, ‘interleukin-2 mediated signaling’) and immunosuppression processes (‘response to interleukin-4’ and ‘TOR signaling’) (**Figure 4A**). The KEGG pathway analysis results revealed TMR was highly enriched in ‘Notch signaling pathway’, ‘base excision repair’, and ‘DNA replication’. (**Figure 4B**). While the heatmaps show a degree of heterogeneity in gene set enrichment scores across the TMR samples, the overall trend in expression of these annotations is upregulated in TMR. The significantly upregulated and downregulated members of each enriched gene set are presented in **Supplementary Table S5-S6**.

**Figure 4.**
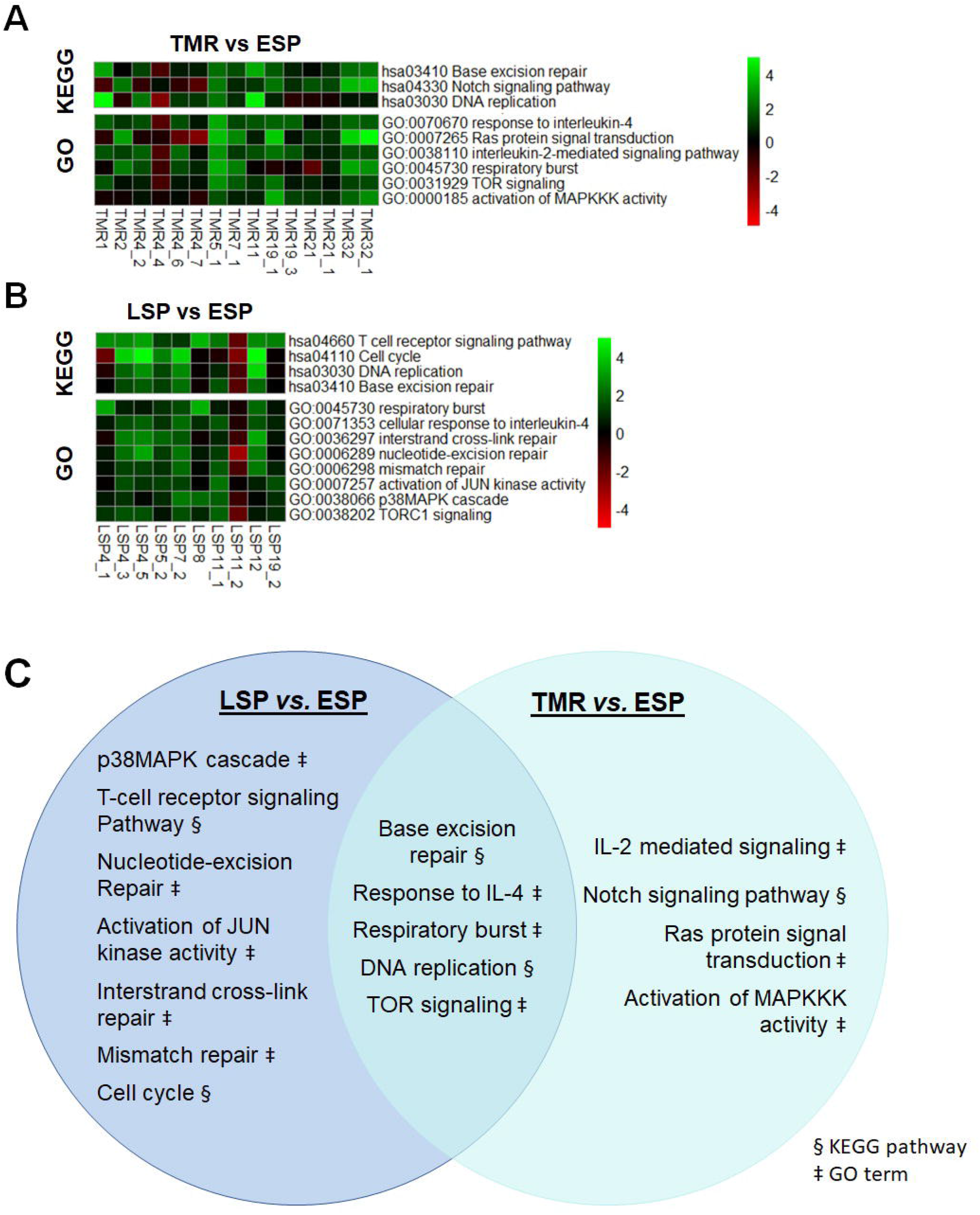
KEGG Pathway and GO enrichment analysis of early (ESP) and advanced staged MF (LSP or TMR) using all available gene expression data. **A.** Heatmaps of select enriched KEGG pathways and GO biological process terms (rows) that are significantly over-represented in TMR vs ESP. **B.** Heatmaps of select enriched KEGG pathways and GO biological process terms (rows) that are significantly over-represented in LSP vs ESP. **C.** Venn diagram showing GO and KEGG annotations common to both LSP and TMR relative to ESP, notably ‘response to interleukin-4’, ‘respiratory burst’, ‘TOR signaling’, ‘DNA replication’ and ‘base excision repair’. KEGG, Kyoto Encyclopedia of Genes and Genomes; GO, gene ontology.

Next, we conducted functional enrichment analyses on LSP vs ESP (reference, **Figure 4B**). Although only a few individual genes were statistically significant between LSP and ESP, GSA identified 29 GO Biological Processes and 14 KEGG pathways overrepresented in LSP (**Supplementary Figure 2**). Among the GO and KEGG hits were five overlapping annotations also captured in the comparison between TMR and ESP (**Figure 4C**). These include three GO terms: ‘respiratory burst’ (inflammatory response), ‘response to interleukin-4’ (Th2 response), and ‘TOR signaling’ (growth and metabolism) as well as two KEGG pathways related to chromatin regulation: ‘DNA replication’ and ‘base excision repair’.

We also identified GO terms and KEGG pathways uniquely enriched in the LSP group compared with ESP (**Figure 4C**), including three DNA damage removal annotations (‘nucleotide-excision repair’ [GO], ‘mismatch repair’ [GO], ‘interstrand cross-link repair’ [GO]), two T-cell function annotations (‘T-cell receptor signaling’ [KEGG], ‘Activation of JUN kinase activity’ [GO]), as well as hits in ‘cell cycle’ [KEGG] and ‘p38MAPK cascade’ [GO]. Enrichment of major DNA repair mechanisms in LSP is consistent with the widespread structural genomic alterations including fusion transcripts, insertions, deletions, and copy number variation described in advanced MF.^10^

Lastly, we performed GSA between TMR and LSP (reference). As expected, expression differences were smaller between LSP and TMR than in any other pairwise comparison, with only four enriched KEGG pathways and seven GO terms (**Supplementary Figure S3**). Compared with LSP, TMR lesions were significantly upregulated in primary metabolic processes including ‘ribosome biogenesis’ ‘biosynthesis of amino acids’, and ‘hexose metabolic process’.

### Other Significant DEGs

Annotation databases like KEGG and GO allow us to collapse complex gene expression data into more manageable biological pathways and functional annotations, but the mapping of genes to pathways is limited by current knowledge of known molecular interaction networks and how gene products interact. To identify potential DEGs that could have been overlooked by functional analysis methods, we manually searched for individual genes that were significantly over-represented in each lesion type to provide a more nuanced understanding of transcriptional networks.

We confirmed overexpression of several genes reported previously in CTCL, such as *KIR3DL2* (>14 fold) and *KIR3DL4* (>6 fold), the members of the killer cell Ig-like receptor family that inhibit natural killer cell-mediated antitumor cytotoxicity. *KIR3DL2* has been implicated in SS^28^ and transformed MF^29^ and the anti-KIR3DL2 monoclonal antibody IPH4102 has recently demonstrated promising clinical activity in patients with refractory CTCL.^30^ LSP and TMR had increased levels of *GNLY* (granulysin, an immune alarmin) and *NCR1* (natural cytotoxicity triggering receptor 1) whose expression was previously found to increase with the progression of MF^31^ and in SS patients with high tumor burden^32^.

Among the 29 DEGs between TMR and LSP, we found a >6 fold decrease in fractalkine Receptor (CX3CR1). CX3CL1 was shown to control effector T cell retention in inflamed skin and in the lungs in atopic dermatitis and asthma^33,34^ and the decrease in its expression may facilitate escapement of malignant cells from the skin to the circulation in late stages MF.

We found ectopic expression of several meiosis-related cancer-testis antigens (meiCT) in TMR lesions. MeiCT antigens are normally present only in germ cells during oocyte development and spermatogenesis and become transcriptionally silent in normal somatic tissues. Ectopic activation of meiCT is a frequent phenomenon in cancer because global deregulation of epigenetic signaling causes unprogrammed gene activation.^35^ While our results confirm upregulated expression of two previously described meiCT antigens (GTSF1 [gametocyte specific factor 1]^36,37^ and STAG3 [Stromal Antigen 3])^38^ we also identified increased expression of seven putative CT antigens previously unreported in CTCL, including CT45A1 (Cancer/Testis Antigen Family 45 Member A1), its paralogue CT45A3 (Cancer/Testis Antigen Family 45 Member A3), BRDT (Cancer/Testis Antigen 9), PRAME (Cancer/Testis Antigen 130), PAGE5 (Cancer/Testis Antigen Family 16 Member 1), SPAG4 (Sperm Associated Antigen 4), and PNMA5 (Paraneoplastic antigen-like 5). These results are in line with the notion that re-expression of a variety of CT antigens in malignant lymphocytes may contribute to increased genomic instability.

### Linking gene mutations to transcriptomic changes

We examined whether the gene mutation pattern in MF could explain some of the observed alterations in the transcriptome. From the previously published catalogue of mutated genes obtained by whole-exome sequencing^17^ we selected the genes which are frequently mutated in TMR and LSP and more expressed in those advanced-stage lesions compared to ESP. We assumed that those genes would be particularly relevant because mutations (especially damaging mutations) cause a compensatory overexpression of the gene.^39^ We found that TMR/LSP lesions showed overexpression of mutated genes representing transcription factors and regulators (*MED12, MYCN, MYC, EGR3, CIC*), genes involved in genome integrity (*ERCC2, POLE*), the elements of the membrane receptor signalling network of integrated JAK-STAT, MEK/ERK, and PI3K/AKT/mTOR pathway (*PLCG1, INPPL1, VAV1, AKT1, PPP2R1A, NFKB2, CARD11, TRAF2, JAK3, STAT3*). Mutations in those genes have been linked to the signaling pathways in different cancers as summarized in **Figure 5A**.

**Figure 5.**
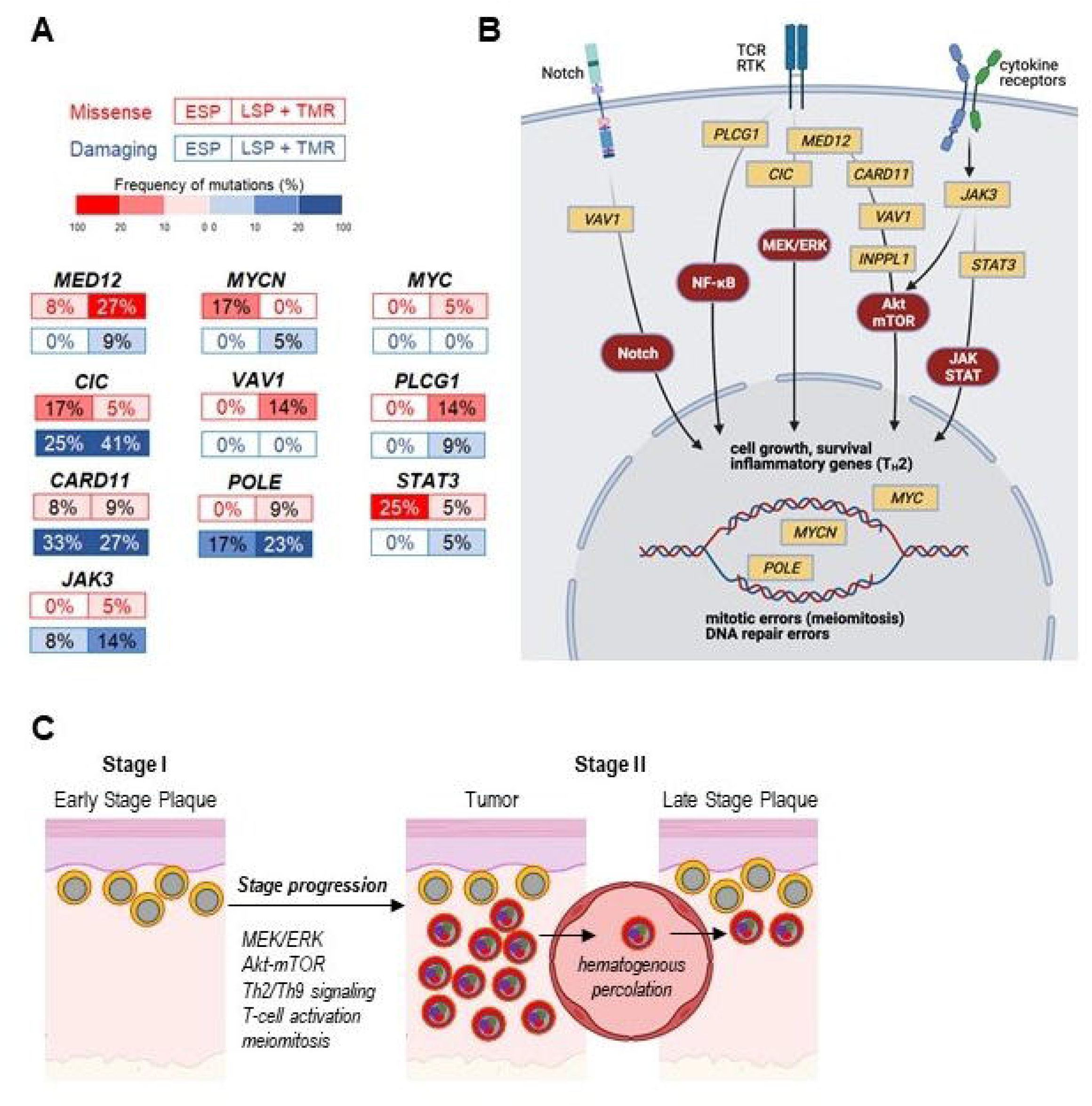
Proposed mechanism of stage progression in MF. **A.** Frequently mutated genes in advanced lesions of MF. The frequencies of non-synonymous mutations (missense or damaging: frameshift mutations, short read insertions and deletions (<6pm), stop gain or stop loss) are shown. Only genes showing compensatory overexpression (significantly increased Log2FC >1, adjusted P <0.05 mRNA levels in TMR vs ESP) are shown. **B.** Possible involvement mutated genes and upregulated signaling pathways during stage progression. The pathways are marked by red ellipses, mutated genes affecting the pathway are plotted in yellow rectangles. **C.** The proposed mechanism of propagation of mutated cells to skin lesions. Early stage plaques contain cells characterized by a low mutation burden^17^ and a characteristic transcriptional signature as shown in *Figure 2*. Accumulation of mutations (colored dots in the “nuclei”) changes the transcriptional signature which enables higher rate of proliferation and enhanced cell survival which lead to the formation of a tumor (high-grade malignant cells marked in red). Those cells can enter the circulation, e.g. due to loss of expression of CX3CR1 and can colonize the already existing plaques (cancer self-seeding) or new areas of the skin in a quasi-metastatic process as proposed by us previously^14^.

## Discussion

Our study addresses the poorly elucidated question of the mechanisms responsible for the development of tumors in MF, an event that defines the progression from the early to the advanced stage of the disease and which is associated with a dramatic increase in mortality. Among the studies where transcriptomic changes during stage progression were captured^37,40–42^ only two reports^36,43^ and a meta-analysis^41^ compared DEGs directly between early and advanced lesions. The novel aspects of this study is that we not only compared skin lesions in early and late-stage patients, but we included different morphological types of MF lesions (the plaques and the tumors: ESP, LSP, TMR) and used cancer cell enriched, microdissected tissue of the high tumor cell fraction (mean purity [SD] 74.6 [20.4]%) which allowed us to largely limit the contribution of transcripts originating from non-neoplastic cells in the skin (epidermis, inflammatory cells, fibroblasts, and other stromal cells).

Compared with early-stage plaques (ESP), the tumor (TMR) samples showed an increased frequency of mutation in genes representing transcription factors and regulators, genes involved in genome integrity, JAK-STAT, MAPK/ERK, and PI3K/AKT/mTOR pathways **(Figure 5A)**. Although detailed mechanistic studies are needed to determine the relative importance of those mutations, overlaying the mutated genes on the known signaling pathway elements reveals their likely relevance in cancer progression. It was shown previously that mutated *PLCG1* increases the survival signaling via NF-kB, NFAT and AP-1.^11,44^ Mutated *STAT3* was also shown to have a pathogenic role in CTCL.^45,46^. Mutations in *CARD11*^9,11^ and *JAK3*^2,47^ are also supposed to increase the growth and survival of malignant T-cells in CTCL and are well-known driver mutations in other hematological malignancies.

Components of MEK/ERK pathway are reproducibly overexpressed in TMR and our mutation data revealed two potentially novel and previously unreported mechanisms of its activation **(Figure 5A)**. *MED12* is a tumor suppressor gene which in normal cells inhibits the transforming growth factor β (TGFβ), Akt/mTOR and MAPK/ERK signaling.^48–50^ Interestingly, loss of function of mutated *MED12* was shown to be the main driver in certain tumors by activating RAS/MAPK/ERK^49–51^. *MED12* was also linked to drug resistance, which might be relevant in CTCL which is notoriously resistant to chemotherapy. Mutations in *Capicua (CIC)* may amplify MAPK/ERK signaling because CIC is a transcriptional repressor counteracting activation of genes downstream of ERK and contributes to the tumorigenesis of different solid tumors.^52–54^

VAV1 is a hematopoietic-specific RHO guanine exchange factor and its loss of function causes T-cell leukemias by increasing the Notch1-dependent ICN1 signaling due to inhibition of ICN1 ubiquitinylation and degradation.^55^ Mutations in VAV1 were reported previously^56^ but were linked to PI-3-kinase signaling rather than Notch signaling. However, previous reports from our lab documenting an increase in Notch1 and its signaling in advanced MF^57^ confirmed by the Notch1 overexpression in TMR seen here, seem to indicate that VAV1-Notch1 axis may play in MF progression.

The notion that MF is a genetically stable tumor and does not hypermutate^58^ has recently been refuted by genome and exome sequencing data showing numerous mutations and subclonal diversification even in early stages.^10,17,59^ The high rate of mutations can be explained in part by our current findings showing dysregulation of major pathways controlling DNA repair mechanisms (nucleotide excision repair, mismatch repair and base excision repair). Although DNA repair mechanisms have not been studied in MF in detail, it is known that mutations affecting DNA repair genes such as *POLE, MYCN*, or *ATM* may contribute to genomic instability in CTCL.^17,60–62^

Another, novel mechanism of genomic instability might be related to abnormal regulation of mitosis. We identified ectopic expressions of five testis-specific genes (*CT45A1, CT45A3, STAG3, GTSF1*, and *REC8*) which are only expressed in testicular germ cells. Ectopic expression of cancer-testis (CT) antigens was previously found in carcinomas of the bladder,^63^ lung,^64^ liver^65^ and suspected to play an important role in maintaining cell survival through inhibition of apoptosis (i.e. down regulating p53 and p21 tumor suppressor genes) and promotion of chromosomal instability (i.e. generation of double-strand breaks leading to loss of heterozygosity and chromosomal arrangements). CT genes (*SYCP1, SYCP3, REC8, SPO11, STAG3, GTSF1*) have previously been reported in advanced CTCL and proposed to reflect the reactivation of meiotic and gametogenic programs in mitotic cells.^37,38^ Our data documenting a dramatic increase in CT expression (10-30 fold up-regulation in MF tumors) support the putative role of meiomitosis^66,67^ as a mechanism of genomic instability in MF.

MF is a spatially discontinuous neoplasm composed of circumscribed lesions of the skin. This calls for a question of whether those lesions evolve in isolation or whether malignant cells percolate between the lesions. Clinical observations support the latter notion; it is known that the development of the tumor in one area of the skin changes the clinical course to be more unfavourable and that those patients are more likely to develop additional tumors. We have therefore hypothesized that high-grade malignant cells in the tumors escape to the circulation and may metastatically seed the skin - either to the already existing lesions or to the new, previously uninvolved areas. The comparisons of global expression profiles of ESP, LSP and TMR seem to confirm our hypothesis. The principal component clustering of transcriptomes showed that LSP and TMR overlapped in two clusters (clusters B1 and B2, Figure 2) and separated well from ESP. The same was previously seen on the mutational level: LSP resembled tumors and differed from the ESP.^17,68^ This shows that stage progression is associated with global genomic and transcriptomic changes in all skin lesions, in spite of the fact that the tumor formation is only present in a discrete skin area. Thus, cancer cells in MF are likely to percolate through the skin, occasionally seeding new areas and the existing lesions, the latter case resembling tumor self-seeding or cell exchange between distant metastases (Figure 5B).^15,16,69,70^ It is conceivable that early eradication of emerging tumors (e.g. by aggressive radiotherapy) would limit hematogenous percolation of malignant cells and retard the colonization of the skin with high-grade mutated cells.

## Acknowledgements

We extend our gratitude to Dr. Thomas Salopek, Mrs. Rachel Doucet, and the nursing staff of Kaye Edmonton Clinic for their assistance with sample collection. This study was supported by grants from the following sources: Canadian Dermatology Foundation (CDF RES0035718), University Hospital Foundation (University of Alberta), Bispebjerg Hospital (Copenhagen, Denmark), Danish Cancer Society (Kræftens Bekæmpelse R124-A7592 Rp12350) and an unrestricted research grant to R.G. from the Department of Medicine, University of Alberta. M.X. was supported by an Alberta Innovates Summer Research Studentship.

## Author’s contributions

M.X. contributed to study design, conducted bioinformatics, analyzed the data, prepared the figures, and wrote the manuscript. RG designed the experiments, supervised data analysis, prepared the figures, and wrote the manuscript. DH and FZ conducted bioinformatics analysis. AI and SO performed wet-lab experiments. AS contributed to writing and editing the manuscript. All authors reviewed all versions of the paper and approved the final version of this manuscript.

## Footnotes

1 The accession number will be provided upon acceptance of the paper. For review purposes the RNA sequencing data can be requested via contacting the corresponding author.

## Notes

**Conflict of Interest:** RG received advisory board honoraria from Kyowa Kirin, Sanofi, Recordati Rare Diseases, and Mallinckrodt and obtained research funding from Sanofi and Sun Pharma.

**Funding:** This study was supported by grants from the following sources: Canadian Dermatology Foundation (CDF RES0035718), University Hospital Foundation (University of Alberta), Bispebjerg Hospital (Copenhagen, Denmark), Danish Cancer Society (Kræftens Bekæmpelse R124-A7592 Rp12350), and an unrestricted research grant to R Gniadecki from the Department of Medicine, University of Alberta. M Xiao was supported by an Alberta Innovates Summer Research Studentship. A Sivanand was supported by scholarships from the Canadian Institutes of Health Research (CIHR), Alberta Innovates, and the University of Alberta. These funding organizations had no role in the design or conduct of the research. **What is known:** - Mycosis fungoides (MF) is the most common cutaneous T-cell lymphoma characterized by a favourable prognosis in the early patch/plaque stage. - Development of tumors heralds progression to the advanced stage and a significant increase in mortality. **What’s new:** - Tumor progression is associated with recurrent mutations which can be linked to the upregulation of signaling pathways controlling cell proliferation, survival, mitosis, and DNA repair. - Percolation of malignant cells between lesions and tumor self-seeding is likely to mediate stage progression in MF. **Translational message:** The development of cutaneous tumors in MF heralds stage progression and increased mortality. MF tumors show upregulation in several targetable signaling pathways involved in cell proliferation and survival. Ectopic expression of cancer-testis genes may explain mitotic aberrations in MF tumors. We also propose that high-grade malignant cells can spread hematogenously and seed other skin lesions (tumor self-seeding). Early, aggressive treatment of tumors may prevent tumor self-seeding and improve patient prognosis.

https://github.com/d-henness/deseq2_project

### Competing Interest Statement

RG received advisory board honoraria from Kyowa Kirin, Sanofi, Recordati Rare Diseases, and Mallinckrodt and obtained research funding from Sanofi and Sun Pharma.

