## Supplementary Figure for "Transcriptomic Changes During Stage Progression of Mycosis Fungoides"

**Supplementary Table S1**: Patient characteristics and samples included in the study

| Patient ID (age [years], sex [M-male, F-female]) | Sample ID[a] | Lesion type | Tumor cell Fraction | Diagnosis and stage at the time of the biopsy | Disease course (observation time, months)[b] | Previous therapy[c] |
|---|---|---|---|---|---|---|
| MF1 (78, F) | MF1 | Tumor | NA | Mycosis Fungoides IIB | SD (26) | RTx*, PUVA, CHOP*, methotrexate* |
| MF2(83, M) | MF2 | Tumor | 0.91309 | Mycosis Fungoides IIB | SD (24) † | TS, RTx*, IFN*, nbUVB |
| MF4 (69, M) | MF4_1P | Plaque | 0.8797 | Mycosis Fungoides IIB | SD (30) | TS, RTx, methotrexate*, surgery, nbUVB |
| | MF4_2T | Tumor | 0.6679 | | | |
| | MF4_3P | Plaque | 0.7337 | | | |
| | MF4_4T | Tumor | 0.91177 | | | |
| | MF4_5P | Plaque | 0.8036 | | | |
| | MF4_6T | Tumor | 0.7237 | | | |
| | MF4_7T | Tumor | 0.8088 | | | |
| MF5 (44, F) | MF5_1T | Tumor | 0.5599 | Folliculotropic Mycosis Fungoides IIB | SD (31) | TS, IFN*, vorinostat*, RTx* |
| | MF5_2P | Plaque | 0.3567 | | | |
| MF7 (62, M) | MF7_1T | Tumor | 0.7128 | Mycosis Fungoides IVA2 | SD (17) † | TS, romidepsin*, RTx*, vorinostat*, IFN*, nvUVB, PUVA* |
| | MF7_2P | Plaque | 0.6203 | | | |
| MF8 (54, F) | MF8P | Plaque | 0.92583 | Folliculotropic Mycosis Fungoides IIIB | SD (36) | TS*, nbUVB*, PUVA*, IFN*, ECP*, CHOEP*, hyper-CVAD*, allogeneic HSCT* |
| MF9 (42, F) | MF9P | Plaque | 0.90173 | Mycosis Fungoides IA | PD IVA (36) | TS, PUVA, CHOEP*, autologous HSCT* |
| MF10 (56, M) | MF10P | Plaque | 0.92182 | Mycosis Fungoides IB | SD (37) | TS, nbUVB* |
| MF11 (56, M) | MF11T | Tumor | 0.8229 | Mycosis Fungoides IIB | SD (37) | TS, nbUVB*, RTx* |
| | MF11_1P | Plaque | 0.5558 | | | |
| | MF11_2P | Plaque | 0.5545 | | | |
| MF12 (66, M) | MF12P | Plaque | 0.91397 | Mycosis Fungoides IVA2 | SD (3) † | TS, methotrexate*, acitretin*, ECP* |
| MF15 (65, M) | MF15P | Plaque | 0.91966 | Mycosis Fungoides IB | SD (36) | TS, nbUVB, allitretinoin* |
| MF19 (74, M) | MF19_1T | Tumor | 0.98573 | | SD (13) † | |

| | | | | | | |
|---|---|---|---|---|---|---|
| | MF19_2P | Plaque | 0.6727 | Mycosis Fungoides IIB | | PUVA, IFN*, MTX*, RTx* |
| | MF19_3T | Tumor | 0.96316 | | | |
| MF20 (70, M) | MF20 | Plaque | 0.90167 | Mycosis Fungoides IB | SD (33) | TS*, PUVA* |
| MF21(71, F) | MF21 | Tumor | NA | Mycosis Fungoides, IVB | SD (33) | Prednisone, cyclosporine, phototherapy, IFN*, isotretinoin*, RTx*, brentuximab*, total body electron beam radiation* |
| | MF21_1 | Tumor | NA | | | |
| MF23_1 (69, F) | MF23_1P | Plaque | 0.2388 | Mycosis Fungoides, IA | PD, IB (30) | TS, IFN*, nbUVB |
| MF25 (48, F) | MF25P | Plaque | 0.8527 | Mycosis Fungoides IB | SD (29) | TS*, nbUVB* |
| MF26 (76, M) | MF26P | Plaque | 0.5049 | Mycosis Fungoides IB | PD IIB (21) | TS, RTx*, IFN* |
| MF27 (71, M) | MF27P | Plaque | 0.92312 | Mycosis Fungoides IA | SD (24) | TS* |
| MF29 | MF29_1P | Plaque | 0.8926 | Mycosis Fungoides IA | PD IIB (15) | TS, nbUVB, surgery*, methotrexate* |
| | MF29_2P | Plaque | 0.92178 | | | |
| MF30 (62, M) | MF30P | Plaque | 0.6289 | Mycosis Fungoides IB | PD IIB (21) | TS*, RTx*, IFN*, PUVA*, cobomarsen |
| MF32(49, M) | MF32T | Tumor | 0.6527 | Mycosis Fungoides IVA | SD (38) | RTx*, IFN*, PUVA*, gemcitabin*, CHOP* |
| | MF32_1T | Tumor | 0.8014 | | | |
| MF43(60, M) | MF43P | Plaque | 0.2114 | Folliculotropic Mycosis Fungoides IA | SD (22) | TS, RTx* |

[a]P, plaque; T, tumor

[b]SD, stable disease (no stage progression during observation time), PD, progressive disease

[c]TS, topical steroid; PUVA, psoralen, and ultraviolet A photochemotherapy; nbUVB, narrow-band UVB; IFN, interferon α, RTx, radiotherapy; ECP ,extracorporeal photopheresis; CHOEP, CHOP, cyclophosphamide, hydroxydaunorubicin, vincristine, prednisone; CVAD, cyclophosphamide, vincristine, doxorubicin, dexamethasone followed by methotrexate and cytarabine.

*therapy administered after biopsy

† deceased

**Supplementary Figure S1.** Clustered heatmap of (A) KEGG pathway and (B) GO BP enrichment analysis in TMR (comparison) vs. ESP (reference, FDR <0.05). Red or green indicates up-regulated or down-regulated gene sets respectively. The legend shows the colour scaling with normalized enrichment values.

# A

#### Enriched KEGG pathways in TMR

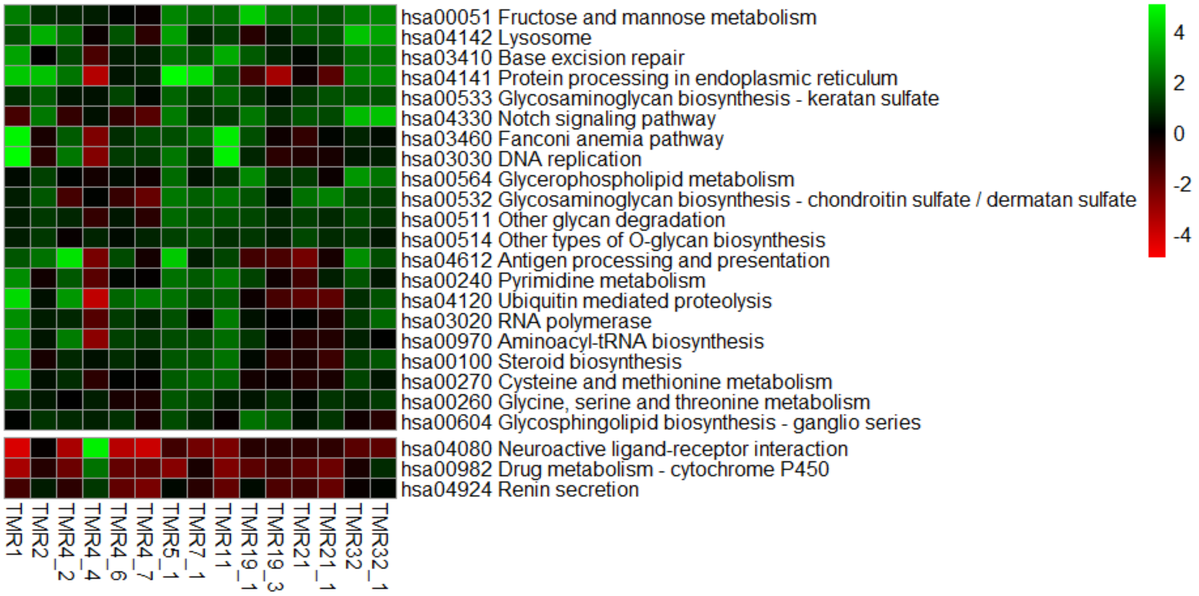

### B Enriched GO terms in TMR

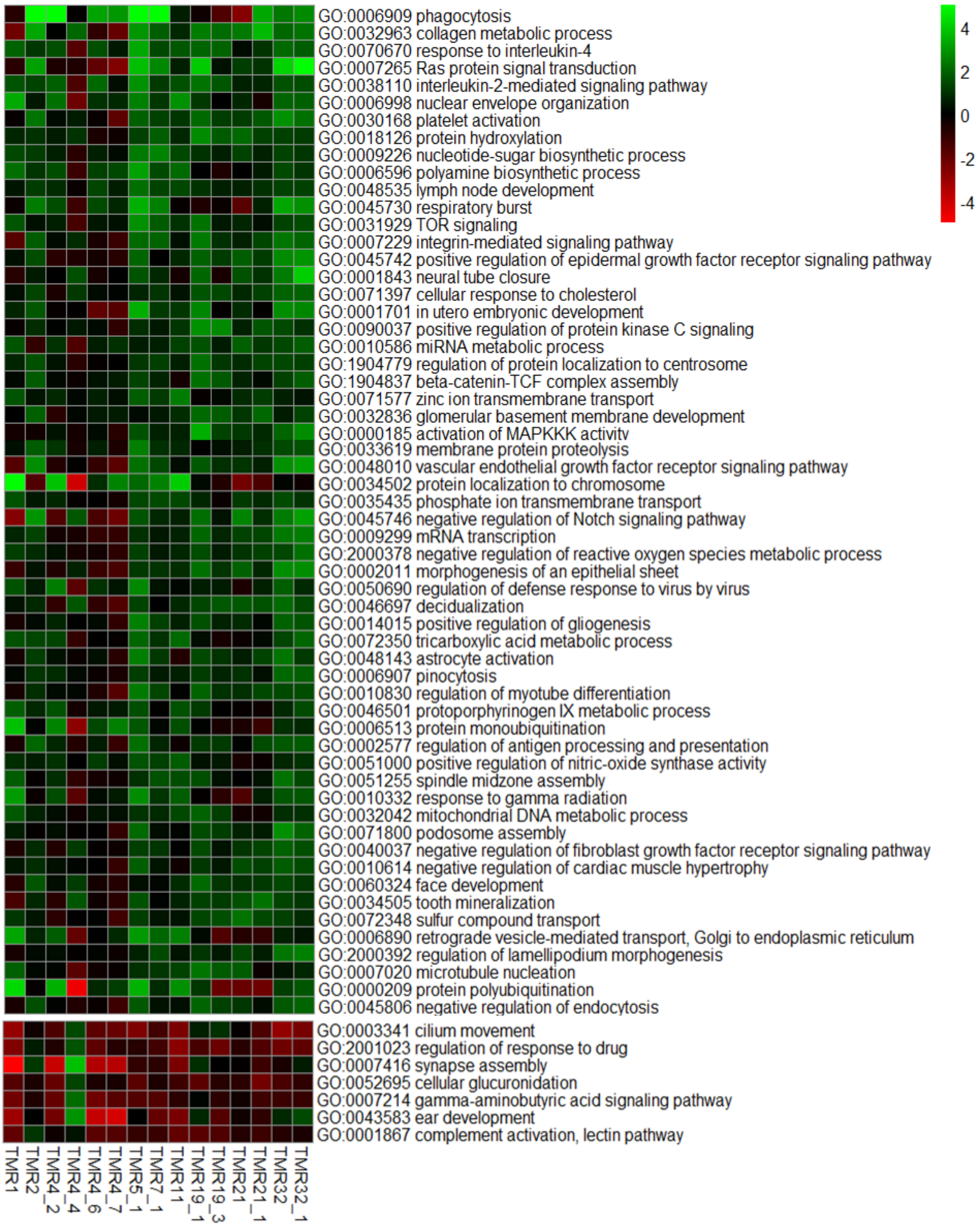

**Supplementary Figure S2.** Clustered heatmap of (A) KEGG and (B) GO BP pathway enrichment analysis in LSP (comparison) vs. ESP (reference, FDR <0.05) The legend shows the colour scaling with normalized enrichment values.

# A

#### Enriched KEGG pathways in LSP

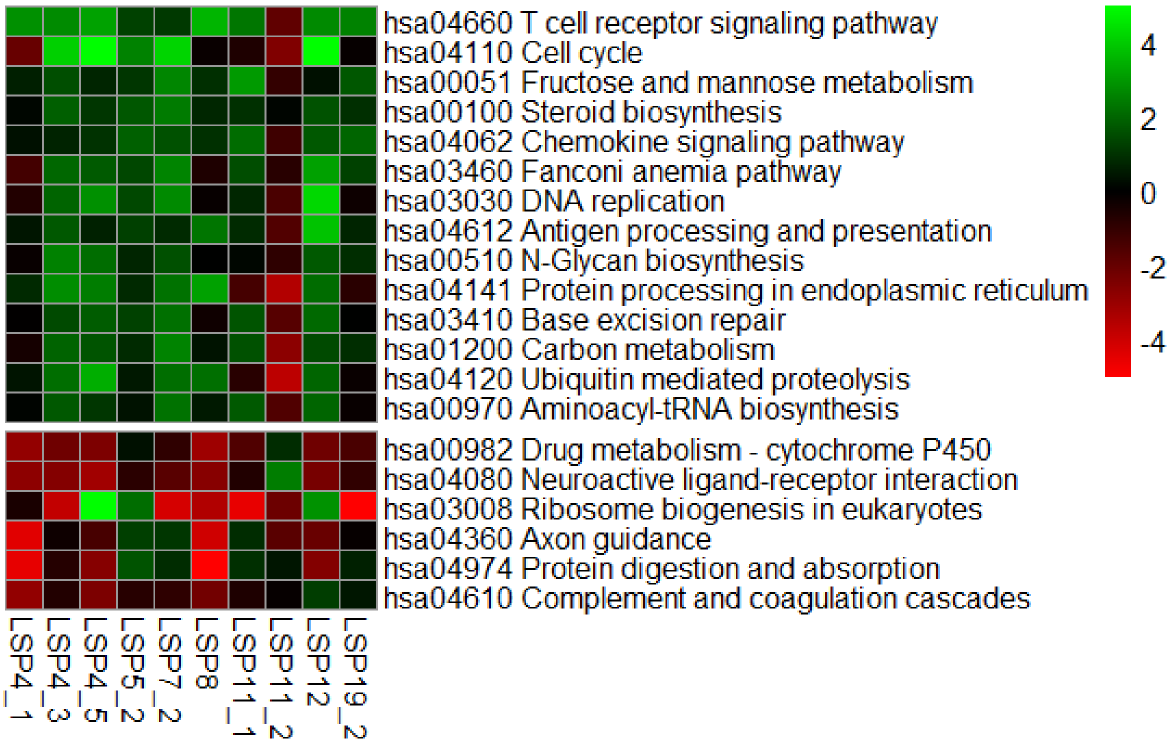

### B Enriched GO terms in LSP

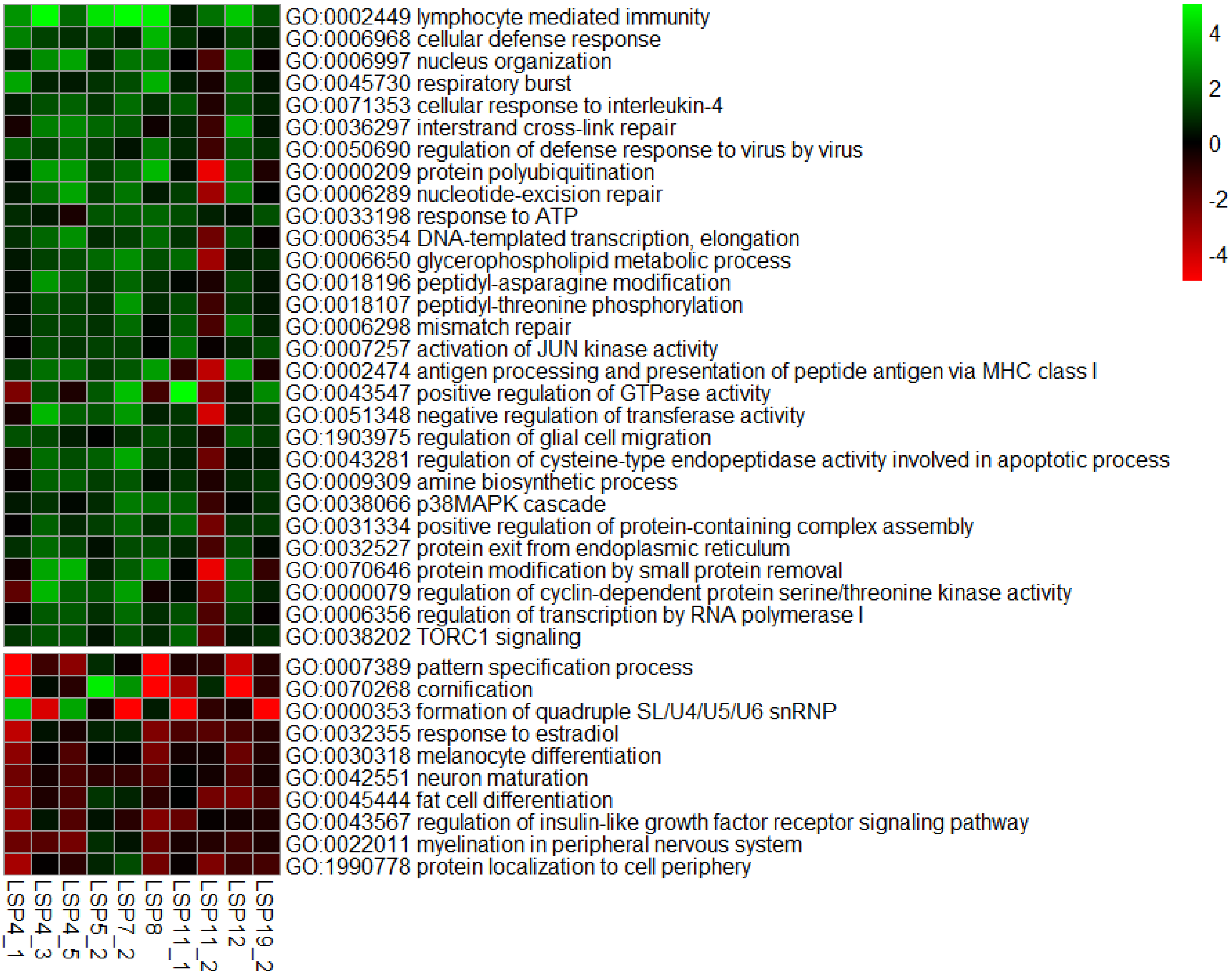

**Supplementary Figure 3**. Heatmaps of select enriched KEGG pathways and GO biological process terms (rows) that are significantly over-represented in TMR (comparison) vs LSP (reference).

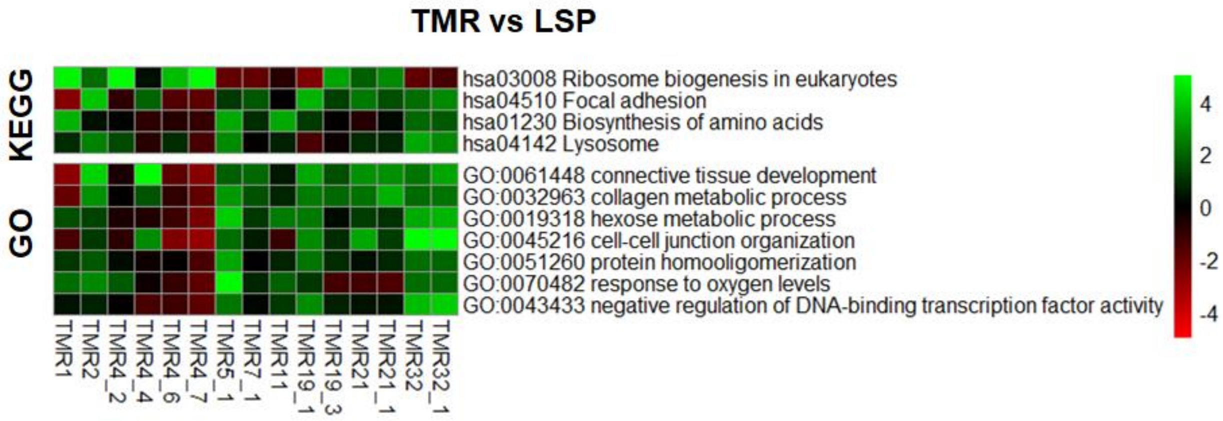

**Supplementary Table S2:** 1,154 differentially expressed genes between TMR (comparison) and ESP (reference).

*For review purposes the full list of 1,154 DEGs can be requested via contacting the corresponding author. An excel sheet will be similarly provided upon acceptance of the paper.*

**Supplementary Table S3**: 26 differentially expressed genes between LSP (comparison) and ESP (reference)

| Gene name | baseMean | log2FoldChange | lfcSE | stat | pvalue | padj |
|---|---|---|---|---|---|---|
| *FSTL5* | 9.026398423 | -6.466891544 | 1.299583005 | -4.976128124 | 6.49E-07 | 0.003515128 |
| *NPY5R* | 3.913757531 | -4.277668283 | 1.032780021 | -4.141896819 | 3.44E-05 | 0.044795801 |
| *AC096644.3* | 8.007872255 | -3.996076043 | 0.935685778 | -4.270745732 | 1.95E-05 | 0.031186851 |
| *LGR5* | 9.373866886 | -3.698040853 | 0.859697198 | -4.301562062 | 1.70E-05 | 0.029021832 |
| *FMO2* | 33.72637371 | -2.913389872 | 0.705236912 | -4.131079675 | 3.61E-05 | 0.045150978 |
| *CRYAB* | 46.74613724 | -2.784145353 | 0.621975933 | -4.476291128 | 7.60E-06 | 0.016462611 |
| *DYNC1I1* | 14.70044277 | -2.152144276 | 0.499754984 | -4.306398824 | 1.66E-05 | 0.029021832 |
| *TCEAL4* | 188.5014654 | -1.897071866 | 0.446235326 | -4.251281234 | 2.13E-05 | 0.031412149 |
| *AC253572.2* | 256.1861672 | 1.994646703 | 0.462939581 | 4.30865449 | 1.64E-05 | 0.029021832 |
| *SLC9A7P1* | 27.72439827 | 2.328172945 | 0.558679262 | 4.167280052 | 3.08E-05 | 0.041759657 |
| *NCR1* | 49.93814809 | 2.561864948 | 0.555048804 | 4.615567006 | 3.92E-06 | 0.011587146 |
| *AC109326.1* | 213.2343279 | 2.72817134 | 0.65128611 | 4.188898396 | 2.80E-05 | 0.039625128 |
| *IL1B* | 63.01621824 | 2.922194361 | 0.649494288 | 4.499184078 | 6.82E-06 | 0.015841906 |
| *GNG4* | 86.12553035 | 3.007565925 | 0.705455769 | 4.263294821 | 2.01E-05 | 0.031186851 |
| *GNLY* | 529.2848515 | 3.495838625 | 0.63834942 | 5.476371584 | 4.34E-08 | 0.000352876 |
| *SCG2* | 63.76591633 | 3.503970256 | 0.753975484 | 4.647326514 | 3.36E-06 | 0.011587146 |
| *AQP9* | 37.59933655 | 3.573353216 | 0.786553019 | 4.543054481 | 5.54E-06 | 0.013866773 |
| *PRSS21* | 16.40235373 | 3.743313324 | 0.690123232 | 5.424123038 | 5.82E-08 | 0.00037871 |
| *EEF1A2* | 70.27665886 | 3.818498111 | 0.824475521 | 4.631426902 | 3.63E-06 | 0.011587146 |
| *U3* | 48.14861482 | 4.08210575 | 0.866332348 | 4.711939663 | 2.45E-06 | 0.009972145 |
| *ITPKA* | 8.548215077 | 4.796408151 | 0.977643574 | 4.906090805 | 9.29E-07 | 0.004315382 |
| *RPPH1* | 17013.96395 | 4.962303366 | 1.114079467 | 4.454173615 | 8.42E-06 | 0.017113407 |
| *LINC01221* | 21.64728762 | 7.211952362 | 1.58155628 | 4.560035235 | 5.11E-06 | 0.013857322 |
| *CU639417.1* | 188.3042615 | 24.61644966 | 1.688285779 | 14.58073625 | 3.72E-48 | 1.21E-43 |
| *RMRP* | 12018.05054 | 29.65066388 | 3.695617061 | 8.023197045 | 1.03E-15 | 1.12E-11 |
| *RN7SK* | 57341.98089 | 30 | 3.494630923 | 8.584597533 | 9.12E-18 | 1.48E-13 |

**Supplementary Table S4:** 29 differentially expressed genes between TMR (comparison) and LSP (reference)

| Gene name | baseMean | log2FoldChange | lfcSE | stat | pvalue | padj |
|---|---|---|---|---|---|---|
| *AC066616.2* | 10.96560468 | -4.983903139 | 1.1330916 | -4.398499767 | 1.09E-05 | 0.018869955 |
| *CXCL5* | 14.33505841 | -3.72885946 | 0.882832298 | -4.223746081 | 2.40E-05 | 0.028939892 |
| *FCN1* | 520.6791704 | -3.652877041 | 0.669223026 | -5.45838517 | 4.80E-08 | 0.001157439 |
| *HEMGN* | 19.05711503 | -3.576631383 | 0.835616825 | -4.28022902 | 1.87E-05 | 0.025917612 |
| *CX3CR1* | 235.8199132 | -2.793663289 | 0.626690529 | -4.457803588 | 8.28E-06 | 0.018869955 |
| *RGS18* | 272.1255246 | -2.751444121 | 0.6664156 | -4.128721061 | 3.65E-05 | 0.035149379 |
| *RGS2* | 332.8577427 | -2.687695093 | 0.528779789 | -5.082824925 | 3.72E-07 | 0.004478897 |
| *IL1B* | 63.01621824 | -2.665368052 | 0.619835179 | -4.300123873 | 1.71E-05 | 0.025700353 |
| *MNDA* | 656.4101558 | -2.280893815 | 0.546983909 | -4.169946828 | 3.05E-05 | 0.031909621 |
| *GLT1D1* | 18.71664661 | -2.187589299 | 0.539446827 | -4.055245462 | 5.01E-05 | 0.0421898 |
| *LINC00877* | 36.47388246 | -1.959809583 | 0.483669881 | -4.051957047 | 5.08E-05 | 0.0421898 |
| *AL365203.2* | 24.50199862 | -1.642706655 | 0.349528263 | -4.699782044 | 2.60E-06 | 0.012973965 |
| *AFMID* | 148.0295969 | 1.022422602 | 0.250122241 | 4.087691678 | 4.36E-05 | 0.040366369 |
| *FGF11* | 46.50756074 | 1.956120856 | 0.466450888 | 4.193626612 | 2.75E-05 | 0.031165783 |
| *RN7SL4P* | 231.8060984 | 2.068383899 | 0.509723659 | 4.057853429 | 4.95E-05 | 0.0421898 |
| *LRRC61* | 46.41287322 | 2.335543785 | 0.521374453 | 4.479589995 | 7.48E-06 | 0.018869955 |
| *GOLGA8Q* | 30.61614066 | 2.649857927 | 0.64149298 | 4.130766836 | 3.62E-05 | 0.035149379 |
| *RN7SL5P* | 130.4737184 | 2.703189094 | 0.64585746 | 4.18542676 | 2.85E-05 | 0.031165783 |
| *DNMT3B* | 101.6056972 | 2.909152717 | 0.62488131 | 4.655528451 | 3.23E-06 | 0.012973965 |
| *NACAD* | 20.4465357 | 3.011063221 | 0.682875133 | 4.409390639 | 1.04E-05 | 0.018869955 |
| *FHDC1* | 102.8264954 | 3.018362986 | 0.606100883 | 4.979967977 | 6.36E-07 | 0.00510645 |
| *FAM30A* | 249.4693492 | 3.036445331 | 0.66555534 | 4.562273259 | 5.06E-06 | 0.017413845 |
| *TMPRSS3* | 82.36691901 | 3.14293059 | 0.708637751 | 4.435172391 | 9.20E-06 | 0.018869955 |
| *AC015818.9* | 7.582609195 | 3.864004852 | 0.824405001 | 4.687022577 | 2.77E-06 | 0.012973965 |
| *PMS2P10* | 14.40323427 | 4.236793228 | 0.991741707 | 4.272073263 | 1.94E-05 | 0.025917612 |
| *QRFPR* | 18.10790204 | 4.284695504 | 0.977755915 | 4.382172931 | 1.18E-05 | 0.018869955 |
| *LRP2* | 17.75579126 | 4.840793655 | 1.088271118 | 4.448150443 | 8.66E-06 | 0.018869955 |
| *CSPG4P13* | 7.947291706 | 5.098997239 | 1.201787536 | 4.242844169 | 2.21E-05 | 0.02798187 |
| *AC012488.2* | 6.50702245 | 6.378346941 | 1.452385516 | 4.391634915 | 1.13E-05 | 0.018869955 |

**Supplementary Table S5.** Kyoto Encyclopedia Analysis of Genes and Genomes pathway analysis of DEGs associated with TMR (comparison) vs ESP (reference)

| KEGG Pathway | Gene name | log2FoldChange | Adjusted P |
|---|---|---|---|
| hsa03410 Base excision repair | *PARP1* | 0.87 | 0.006 |
| | *POLE* | 1.18 | 0.008 |
| | *POLD4* | 1.16 | 0.019 |
| | *POLD1* | 1.59 | 0.024 |
| | *OGG1* | 0.69 | 0.025 |
| | *XRCC1* | 1.14 | 0.040 |
| hsa04330 Notch signaling pathway | *DTX3* | 1.4 | 0.015 |
| | *MFNG* | 1.2 | 0.016 |
| | *DVL2* | 1.07 | 0.026 |
| | *CIR1* | -0.82 | 0.031 |
| | *DVL3* | 1.06 | 0.043 |
| | *NOTCH1* | 1.01 | 0.054 |
| hsa03030 DNA replication | *POLE* | 1.18 | 0.008 |
| | *POLD4* | 1.16 | 0.019 |
| | *POLD1* | 1.59 | 0.024 |

**Supplementary Table S6.** Gene Ontology analysis of DEGs associated with TMR (comparison) vs ESP (reference)

| Gene Ontology | Gene name | log2FoldChange | Adjusted P |
|---|---|---|---|
| GO:0070670 response to interleukin-4 | *JAK3* | 1.33 | 0.018 |
| | *CORO1A* | 1.33 | 0.018 |
| | *STAT6* | 0.96 | 0.027 |
| | *ADAMTS13* | 1.61 | 0.034 |
| | *IL2RG* | 1.47 | 0.037 |
| | *XBP1* | 1.42 | 0.039 |
| | *IL4R* | 1.13 | 0.041 |
| GO:0038110 interleukin-2-mediated signaling pathway | *IL2RB* | 2.64 | <0.001 |
| | *JAK3* | 1.33 | 0.018 |
| | *SHC1* | 0.87 | 0.026 |
| | *STAT5A* | 0.96 | 0.030 |
| | *IL2RG* | 1.47 | 0.037 |
| | *PTK2B* | 0.95 | 0.041 |
| GO:0045730 respiratory burst | *CYBC1* | 0.99 | 0.006 |
| | *JCHAIN* | 2.83 | 0.012 |
| | *PIK3CD* | 1.19 | 0.041 |
| | *CYBA* | 1.14 | 0.043 |
| GO:0031929 TOR signaling | *AKT1* | 1.29 | 0.012 |
| | *DGKQ* | 1.42 | 0.012 |
| | *CCDC88A* | -0.92 | 0.016 |
| | *GSK3A* | 1.36 | 0.017 |
| | *CASTOR2* | 1.21 | 0.019 |
| | *CASTOR1* | 1.57 | 0.023 |
| | *ARAF* | 0.95 | 0.024 |
| | *KPTN* | 1.72 | 0.026 |
| | *SPAAR* | -1.83 | 0.038 |
| | *CARD11* | 1.45 | 0.044 |
| | *MEAK7* | 1.18 | 0.048 |
| GO:0000185 activation of MAPKKK activity | *MAP4K1* | 1.54 | 0.011 |
| | *GADD45G* | 2.15 | 0.040 |
| GO:0007265 Ras protein signal transduction | *LIMK1* | 1.51 | 0.002 |
| | *RAB40C* | 1.85 | 0.002 |
| | *RAC3* | 2.55 | 0.003 |
| | *GDI1* | 1.69 | 0.003 |
| | *LAT* | 1.69 | 0.004 |
| | *ARRB1* | 1.06 | 0.006 |

| | | | |
|---|---|---|---|
| | *RASSF1* | 1.07 | 0.007 |
| | *GPR35* | 2.2 | 0.010 |
| | *ARHGAP1* | 1.37 | 0.012 |
| | *RHOT2* | 1.69 | 0.013 |
| | *CYTH2* | 1.21 | 0.016 |
| | *KSR1* | 1.28 | 0.022 |
| | *ARHGDIA* | 1.42 | 0.024 |
| | *CELSR1* | 1.88 | 0.024 |
| | *SHC1* | 0.87 | 0.026 |
| | *CSF1* | 1.55 | 0.030 |
| | *EPS8L1* | 1.91 | 0.030 |
| | *RANGRF* | 0.94 | 0.032 |
| | *FBP1* | 1.62 | 0.033 |
| | *MYO9B* | 1.16 | 0.035 |
| | *RASAL3* | 1.45 | 0.036 |
| | *RFXANK* | 1.16 | 0.037 |
| | *RALGDS* | 1.36 | 0.037 |
| | *MADD* | 0.81 | 0.037 |
| | *DENND4B* | 0.89 | 0.039 |
| | *OBSCN* | 1.42 | 0.039 |
| | *ABR* | 1.19 | 0.040 |
| | *MAPK11* | 1.45 | 0.041 |
| | *VAV2* | 1.26 | 0.041 |
| | *SGSM3* | 1.20 | 0.043 |
| | *VAV1* | 0.99 | 0.050 |
